# Oxygen consumption of drift-feeding rainbow trout: the energetic tradeoff between locomotion and feeding in flow

**DOI:** 10.1101/2019.12.26.889055

**Authors:** J.L. Johansen, O. Akanyeti, J.C. Liao

**Affiliations:** The Whitney Laboratory for Marine Bioscience, Department of Biology, University of Florida, 9505 Oceanshore Blvd, St Augustine, FL 32080, USA; Hawaii Institute of Marine Biology, University of Hawaii Manoa, HI 96744, USA; Department of Computer Science, Aberystwyth University, Penglais Campus, Aberystwyth SY23 3FL, UK

**Keywords:** energetics, swimming, feeding, flow refuging, Kármán gait, prey capture, turbulence, respirometry

## Abstract

To forage in fast, turbulent flow environments where prey are abundant, predatory fishes must deal with the high associated costs of locomotion. Prevailing theory suggests that many species exploit hydrodynamic refuges to minimize the cost of locomotion while foraging. Here we challenge this theory based on direct oxygen consumption measurements of drift-feeding trout (*Oncorhynchus mykiss)* foraging in the freestream and from behind a flow refuge at velocities up to 100 cm s^-1^. We demonstrate that refuging is not energetically beneficial when foraging in fast flows due to a high attack cost and low prey capture success associated with leaving a station-holding refuge to intercept prey. By integrating optimum foraging theory with empirical data from respirometry and video imaging, we develop a mathematical model to predict when drift-feeding fishes should exploit or avoid refuges based on prey density, size and flow velocity. Our foraging and refuging model provides new mechanistic insights into the locomotor costs, habitat use, and prey selection of fishes foraging in current-swept habitats.

## Introduction

The mechanisms underlying how animals distribute themselves in nature are a topic of major interest to ecologists. One major tenet is that animals seek to maximize their fitness by optimizing the ratio of energy intake to energy usage. In current-swept environments, fishes distribute themselves by exploiting flow refuges and vortices to reduce the cost of locomotion (Cotel et al 2006, Liao et al 2003a, Johansen et al 2008, Wilkes et al 2017). These habitats are also often associated with high prey densities (Hill & Grossman 1993, Hayes et al 2007, Jenkins & Keeley 2010). Theoretical cost-benefit models assert that the distribution pattern of fishes reflect their attempt to minimize energy used for swimming while maximizing their energy intake through foraging (Kiflawi & Genin 1997, Rosenfeld et al 2014, Piccolo et al 2014). However, since empirical data on the cost of foraging are non-existent, these models are prone to prediction inaccuracies.

Here, we develop for the first time a data-driven energetics model that predicts foraging strategy and prey size of fishes as a function of flow velocity. This approach enabled us to address previously inaccessible questions such as, 1) what is the ability of refuging fish to detect and capture prey? (Rosenfeld et al 2014); 2), what are the direct energetic costs of leaving a refuge to capture prey? (Guensch et al 2001); and 3) what is the optimum foraging and refuging strategy? We test the long-standing hypothesis that foraging fish expend less energy when refuging compared to swimming in the freestream. By directly measuring oxygen costs of foraging fishes, we demonstrate that in fast flow foraging fishes do not gain an energetic benefit when refuging.

## Materials and Methods

### Animals

We selected rainbow trout *Oncorhynchus mykiss* (Walbaum) as a common, representative drift-feeding species. Fish were obtained from the Chattahoochee Forest National Fish Hatchery, Georgia, and the Cantrell Creek Trout Farm, North Carolina, USA. Fish were kept in two 473 L circular freshwater tanks maintained at 15 ± 0.5°C (mean ± s.e.m.) with a DS-4-TXV Delta Star Chiller (Aqua Logic Inc, San Diego CA, USA). The fish were kept on a 12:12 light:dark cycle and fed commercial trout pellets daily for a minimum of one week prior to experimentation (Pentair -Dense Culture F2A Pellets).

Trout were divided among two experimental treatments: 14 large trout (total length = 33.0 ± 0.6 cm, weight = 423.8 ± 19.2 g, mean ± s.e.m.) were used to measure oxygen consumption and energy expenditure while swimming, which requires a relatively small volume of water to fish size for accuracy (generally less than 300:1 by volume) (Clark et al 2013, Svendsen et al 2016). 18 smaller trout (total length = 16.3 ± 0.2 cm, weight = 70.3 ± 4.3 g, mean ± s.e.m.) were used to measure prey detection and capture success. For these experiments, it was important that fish had enough space in the experimental flow tank to maneuver and capture drifting prey unconstrained (see below). Trials were conducted within the 12hr daily light regime in order to match the diurnal activities of the study species, and all test subjects were starved for 48hrs before the start of experiments to maximize feeding motivation and to ensure a post-absorptive state for swimming (Niimi & Beamish, 1974).

### Experimental setup

All experiments were conducted using a customized 175L recirculating flow tunnel respirometer (Loligo Systems, Denmark) with a working section of 25 × 26 × 87 cm (width × depth × length). Flow within the working section of the respirometer was calibrated from 0 to 145 ± 0.5 cm s^-1^ (mean ± s.e.m.) using digital particle image velocimetry (DPIV, 5-watt argon-Ion continuous laser, LaVision software). Water within the system was filtered, fully aerated and maintained at a temperature of 15 ± 0.1°C (mean ± s.e.m.) using a thermostat (Auber Instruments TD-100A) attached to a chiller (Aqua Logic DS-4). A D-section cylinder (5 cm diameter) placed in the front of the working section allowed for the creation of a distinct flow refuge within the sealed respirometer (Liao et al 2003b). A Phantom V12 high speed video camera (1024 × 1024, 150 frames per second, Vision Research) was aimed at a mirror angled at 45° below the working section to record the swimming kinematics and feeding behavior of individual trout. In all trials solid blocking effects of the fish in the working section were corrected following Bell & Terhune, 1970 and never exceeded 5%.

### Direct oxygen consumption measurements reveal the energetic cost of locomotion

Individuals were introduced into the respirometer and left to acclimatize overnight (10-12 hrs) at a current velocity of 0.5 L s^-1^, where L is the total body length (i.e. 16.5 cm s^-1^), until oxygen consumption had reduced to a steady state level and the fish had settled into a continuous swimming rhythm. The trial was then started and oxygen consumption was measured at increasing current velocities from 0.5 to 3.0 L s^-1^, at 0.5 increments (i.e. 16.5 – 100.0 cm s^-1^). Oxygen consumption at each velocity was determined over three, consecutive 16-minute measurement cycles. The treatment order in which fish were tested (swimming with no cylinder or refuging behind a cylinder) was randomized. We defined the maximum current velocity as the velocity when fish could no longer hold position and were swept onto the downstream grid for longer than 5 seconds. The experiments were then stopped and the current velocity returned to 16.5 cm s^-1^. The cylinder was then either removed or added depending on the previous treatment, and the fish was left to recover overnight. The experimental protocol was repeated the next morning. After data were collected for each fish both refuging and swimming in the freestream, the trial was ended and the fish returned to its holding tank.

For every oxygen measurement cycle, a 300 s flush, 60 s equilibration and 600 s measurement period was applied following the intermittent flow respirometry methodology of Steffensen et al. (1984) and Steffensen (1989). The flushing period ensured the oxygen concentration throughout the trial did not decrease below 85% of air saturation and avoided any CO_2_ build up. Oxygen levels within the swimming respirometer were measured using a D901 miniature galvanic dissolved oxygen probe (Qubit Systems, Kingston, Canada) and monitored with Autoresp v.1.6 software (Loligo Systems). To reduce bacterial growth and respiration within the system, the respirometer was regularly treated with a Clorox solution and thoroughly flushed with freshwater. This procedure ensured background respiration remained below 3% of the oxygen consumed by each fish during swimming trials, which was subtracted from the overall oxygen consumption of the subject. Specifically, after the fish had been returned to its holding tank, the respirometer was run for two additional 16minute measuring cycles at 33 cm s^-1^ during which the reduction in oxygen saturation in the empty respirometer was measured.

### The energetic cost of attack

We modified our respirometry protocol using the same flush, equilibrium and measuring periods as above. However, to avoid significant increases in oxygen consumption due to digestion (i.e. specific dynamic action, SDA; Alsop & Wood 1997), a 2 mm artificial food particle (herein referred to as the lure) was constructed from synthetic yarn wrapped onto a size 20 fishing hook with the hook point cut off at the bend. The lure allowed for the examination of the energetics of foraging attempts without the interference associated with the cost of digestion. Thin fluorocarbon line (2 lb test) was tied to the lure so that it could be introduced into the respirometer through small access ports located upstream of the cylinder. This allowed repeated trials of drifting the lure naturally with the current towards the test subject. To ensure the lure smelled like food, prior to each experiment it was soaked for 1 hr in 200 ml sterilized water containing 30 commercial trout food pellets (Pentair -Dense Culture F2A Pellets).

At the beginning of a trial, each fish was randomly assigned to an initial “no-cylinder” or “cylinder” treatment. The fish was then introduced into the respirometer and left to acclimatize overnight at a current velocity of 16.5 cm s^-1^ until oxygen consumption had reached a steady state level and the fish had settled into a continuous swimming rhythm (∼ 12 hrs). Current velocity was then gradually increased to the experimental velocity of 68.0 cm s^-1^ at a rate of 0.2 cms^-2^. The fish was left to acclimate for 2-4 hrs at this speed until oxygen consumption rate stabilized.

Feeding responses were stimulated by introducing water scented with food into the flow tank through 1.0 cm diameter access ports. After one minute, the tethered lure was randomly injected into the flow tank through one of three access ports and allowed to drift passively in the freestream flow towards the fish. This procedure enticed the fish to attack the lure, and we recorded the number of attacks as well as the number of times that the fish ignored the lure. Whenever the lure was successfully captured or had passed the fish, it was immediately retracted back to the point of entry by swiftly pulling on the line. After 10 s the lure was re-introduced randomly into the flow tank. We repeated this procedure throughout the 600 s oxygen consumption measuring period, which allowed fish to perform attacks *ad libetum* (n = 14 fish). We were able to measure oxygen consumption across a wide range of attacks because of the inherent variability in behaviors across individuals. Note that we measured cost of attack at one flow speed, and assumed that it increases proportionally with speed.

### Measuring probability of prey capture success

Energy intake trials did not rely on respirometry and could therefore take advantage of real food particles. Prior to these feeding trials, 30 small 3 mm trout food pellets (Pentair -Dense Culture F2A) were soaked in 200 ml sterilized water for an hour and then individually sectioned in two to minimize satiation of the fish during the trial. Small 3 ml pipettes were then filled with either scented water containing no food or with scented water containing food (hereafter referred to as a food particle). All samples of scented water and food particles were then kept in a temperature-controlled bath, until they were used within 1 hr of preparation. Capture success was defined as the number of captured prey divided by the number of attacks.

Experimental treatment and acclimatization protocols were similar to those of the respirometer trials above. Briefly, the fish was introduced into the flow tank and left to acclimatize for 2-4 hrs until it had settled into a continuous swimming rhythm (flow velocity = 16.5 cm s^-1^). The flow velocity was then slowly increased at a rate of 0.2 cms^-2^ to one of five experimental velocities (16.5, 33, 51, 68, or 84 cm s^-1^). Once one of these velocities was reached, fish were left to acclimate for another 5 min. After one minute of exposure to food scented water, the first food particle was injected into the flow tank. For each food particle injected, we recorded whether the fish showed 1) no reaction or 2) attempted to capture the particle, defined as a clear change in direction towards the particle. Capture movements were further divided into successful and failed attempts. These capture behaviors (ignore, successful, failed) were recorded until a total of 5 successful captures were observed. At this point the feeding trial was paused and another current velocity randomly selected. The trial was repeated until each fish had been examined at each of the five experimental flow velocities. This procedure allowed each fish to consume a maximum of 25 food particles, which was equivalent to less than 1% of the fish body weight.

### Experimental Data analysis

The energetic cost of locomotion (for both cylinder refuging and freestream swimming) were plotted as oxygen consumption (mg O_2_ kg^-1^ h^-1^, MO_2_) versus swimming speed (cm s^-1^) and fitted with a three-parameter non-linear power function (y = a + bc^x^, where a + b is the Y-axis intercept) following Roche et al. (2013). Differences in oxygen consumption between groups were analyzed using repeated measures ANOVA and swimming speed as a fixed factor. The energetic cost of attack (for both refuging and freestream swimming) was presented as oxygen consumption of individuals plotted against attack rate per hour, where an attack is defined as a sudden change in direction of the head to intercept the lure. A linear regression was fitted to these data and the slope was used to derive the cost of attacking a single drifting food particle. The difference in slopes was compared using an Analysis of Covariance (ANCOVA), with refuging behavior and current velocity as categorical and continuous variables, respectively. We converted energetic cost values from oxygen consumption to Joules by using a conversion rate of 1 mg O_2_ = 13.56 Joule (Elliott & Davison 1975).

Prey detection was defined as the proportion of food particles that a fish attempted to capture, and was fitted with a best-fit quadratic polynomial curve (y = ax^2^ + bx +c, where c is the Y-axis intercept). Differences in prey detection rate between refuging and freestream swimming fish were compared using a two-way ANOVA with refuge/freestream swimming and current velocity as fixed factors, followed by a Tukey HSD posthoc test to identify differences between groups.

Prey capture success was defined as the proportion of food particles that was successfully ingested. Capture success was plotted against flow velocity and fitted with a best-fit 3-parameter sigmoidal curve (y = a / (1 + exp^-(X-X0)/b^), where a, b and X_0_ are constants). Differences in capture success between refuging and freestream swimming fish were evaluated using a two-way ANOVA with refuge/freestream swimming and flow velocity as fixed factors, followed by a posthoc planned comparison for specific differences between groups and corrected for type 1 errors using False Detection Rate (Benjamini & Hochberg 1995).

### Cost-Benefit Model

We developed a mathematical model to estimate the conditions that would provide the greatest net energy gain (E_Net_) in drift feeding fishes, focusing specifically on: a) optimum flow velocity; b) optimum use of refuges; and c) required prey size and density versus flow velocity. Estimates of *E*_*Net*_ of foraging fish in flow streams were based on gained (*E*_*Gain*_) and lost (*E*_*Cost*_) energy over 1 hr periods,

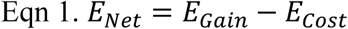

We defined *E*_*Gain*_ and *E*_*Cost*_ as

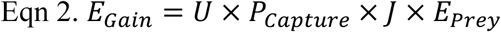

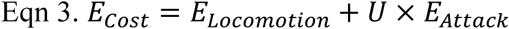

where *U* is the number of attacks and *P*_*Capture*_ is the probability of prey capture success. *E*_*Prey*_ is the energy content of an invertebrate prey (Joule) which is described as a function of prey size after (Jenkins & Keeley 2010),

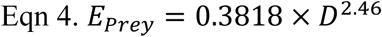

where *D* is the diameter of prey (in mm). *J* is a constant to describe percentage of prey energy available to fish after accounting for digestion and excretion (0.68, Hill & Grossman 1993). *E*_*Locomotion*_ and *E*_*Attack*_ are the energetics cost of swimming and attacking prey, respectively. See Table 1 and 2 for a complete description of model parameters, formulae and data acquisition.

**Table 1.** Overview of model factors and description, data values and sources.

| # | Cost-benefit factors | Definition | Value or method of computation | Source |
| --- | --- | --- | --- | --- |
| A | Prey size distribution | Natural size distribution of invertebrate drift measured in-situ in typical trout streams. Recorded as proportion of whole (%) and diameter size in mm [D] | 51.0%; 0–2 mm<br>43.0%; 2–4 mm<br>5.0%; 4–6 mm<br>0.9%; 6–8 mm<br>0.1%, 8–10 mm | Hughes & Dill 1990<br>Guensch et al 2001<br>Hughes et al 2003 |
| B | Prey density | Number of prey individuals per m<sup>3</sup> in a typical trout stream | 0 - 20 individuals / m<sup>3</sup> | LaPerriere 1981, 1983<br>Hughes & Dill 1990<br>Guensch et al 2001<br>Jenkins & Keeley 2010 |
| C | Foraging area | The total area around an individual fish where it will attack prey | 1.06 m<sup>2</sup> | Hughes et al 2003 |
| D | Diet size composition | The diameter size of prey preferentially consumed | 2 - 10 mm | Hill & Grossman 1993<br>Braaten et al 1997<br>Nakano et al 1999<br>Guensch et al 2001<br>Hughes et al 2003 |
| E | Relative daily food intake | Maximum total daily food intake relative to body size of trout (%) | 3.8 - 5.0% | Brett & Groves 1979<br>Boujard & Medale 1994<br>Nakano et al 1999 |
| F | Daily feeding hours | The number of hours spend feeding each day by a visual predator like trout | Daytime: constant feeding (12 hrs)<br>Night time: resting | Hughes & Dill 1990<br>Hill & Grossman 1993<br>Braaten et al 1997 |
| G | Prey capture time | The time required to capture a prey, including initiation of prey approach, prey intercept and return to starting location (in seconds) | 2 - 20 s | Backman 1984<br>Puckett & Dill 1984<br>Bannon & Ringler 1986 |
|  |  |  |  | Hughes & Kelly 1996<br>Jenkins & Keeley 2010<br>Watz & Piccolo 2011 |
| H | Attack rates | Typical number of preys attacked per hour | 0 - 300 | Hughes & Kelly 1996<br>Hughes et al 2003 |
| I | Energy conversion | Oxygen consumption (mg O<sub>2</sub> kg<sup>-1</sup> hr<sup>-1</sup>) to Joule conversion | 1.00 mg O<sub>2</sub> = 13.56 Joule | Elliott & Davison 1975 |
| J | Prey assimilation efficiency | Energy assimilated of typical invertebrate prey after accounting for digestion and excretion | 58-72% | Elliott 1976<br>Brett & Groves 1979<br>Ware 1982<br>Hill & Grossman 1993<br>Rosenfeld & Boss 2001<br>Jenkins & Keeley 2010 |
| K | Prey mass | Mass (mg) of each prey relative to size (diameter, mm) | $\exp^{(-5.021 + 2.88 \ln(D))}$ | Smock 1980 |
| L | Prey energy content (E<sub>prey</sub>) | Total Joule of each prey item relative to size (diameter in mm) | $0.3818 * D^{2.46}$ | Jenkins & Keeley 2010 |
| M | Flow velocity | Flow velocities typically encountered in streams | 0 – 100 cm s<sup>-1</sup> | Hughes & Dill 1990<br>Gido et al 2000<br>Guensch et al 2001<br>Enders et al 2005<br>Hayes et al 2007<br>Jenkins & Keeley 2010<br>Urabe et al 2011 |
| N | Flow reference velocity | Flow velocity used for measuring the cost of prey attack [R] | 68 cm s<sup>-1</sup> | This study |
| O | Predator mass | Mass of drift-feeding predatory fish | 500 g | Robins & Ray 1986 |

**Table 2.** Model factors, formulae and data sources.

| # | Cost-benefit factor | Formulae | Source |
| --- | --- | --- | --- |
| P | Cost of swimming (mg O<sub>2</sub> kg<sup>-1</sup> hr<sup>-1</sup>) (E<sub>Locomotion</sub>) | $y_{\text{refuge}} = 66.60 + 0.30 * (1.07^M)$<br>$y_{\text{freestream}} = 47.30 + 17.79 * (1.03^M)$ | Empirical data from this study |
| Q | Prey capture success (P<sub>Capture</sub>) | $y_{\text{refuge}} = 1 / (1 + \exp^{-(M-58.811)/-10.579})$<br>$y_{\text{freestream}} = 1 / (1 + \exp^{-(M-86.383)/-16.515})$ | |
| R | Cost of each prey attack (mg O<sub>2</sub> kg<sup>-1</sup> attack<sup>-1</sup>) at flow reference velocity of 68 cm s<sup>-1</sup> (E<sub>Attack</sub>) | Refuge = 1.28 ± 0.18<br>Freestream = 0.78 ± 0.20 |  |
| S | Number of prey available per hour | Prey size distribution * Prey density * Foraging area * Flow velocity<br>[= A * B * C * M] | Derived data |
| T | Maximum food intake per hour | Relative daily food intake * Predator mass / Daily feeding hours<br>[= E * O / F] |  |
| U | Number of attacks per hour | Defined as the lowest value of the following parameters:<br>a) Prey captures per hour [= 3600 / G]<br>b) Number of prey available per hour [S]<br>c) Maximum prey intake per hour [= T / K] |  |
| W | Cost of attacks per hour (mg O<sub>2</sub> kg<sup>-1</sup> hr<sup>-1</sup>) | Cost of each prey attack * Number of attacks per hour * Flow velocity / Flow reference velocity [= R * S * M / N] | Main model |
| X | Total energy cost per hour (E<sub>Cost</sub>) | Cost of swimming per hour + Cost of attacks per hour [= P + W] |  |
| Y | Total energy intake per hour (E<sub>Gain</sub>) | Number of attacks per hour * Prey capture success * Prey energy content * Prey assimilation efficiency [= U * Q * L * J] |  |
| Z | Net energy gain per hour (E<sub>Net</sub>) | E<sub>Gain</sub> - E<sub>Cost</sub> [= Y - X] |  |

### Cost-Benefit Model Analysis

First, we used the cost-benefit model to calculate net energy gain (for both refuging and freestream swimming) over a range of flow velocities (2.0 cm s^-1^ – 100 cm s^-1^, in 1.0 cm s^-1^ increments) and number of attacks (1 – *U*_*max*_). For each experiment, *U*_*max*_ changed dynamically depending on prey availability and eating capacity of fish. Note that the maximum number for *U*_*max*_ per hour was 720 as we assume that each prey attack lasted 5 s (Backman 1984). Prey availability was calculated as

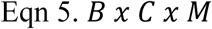

where B is a mean prey density of 20 individuals m^-3^ (Jenkins & Keeley 2010), C is a fish foraging area of 1.06 m^2^ (Hughes et al 2003), and M is flow velocity in cm s^-1^. We assume that consumption capacity of the fish was proportional to its own weight (5% daily, Boujard & Medale 1994) and the weight of prey defined as

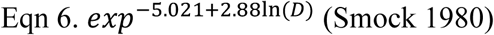

For a given prey density, we used a more realistic non-uniform prey size distribution (0-2 mm (51%), 2-4 mm (43%), 4-6 mm (5%), 6-8 mm (0.9%) and 8-10 mm (0.1%)) in diameter as measured *in-situ* in typical trout streams (Guensch et al 2001).

Since fish in nature do not feed continuously throughout the day, we then sought to identify an optimum strategy when feeding and locomotor behaviours were combined. To do this, we divided a 24 hr day into 12 hr halves, roughly corresponding to night and day and assumed that fish forage exclusively during the day. This allowed us to include three foraging scenarios in our model, 1) foraging while refuging; 2) foraging while freestream swimming; and 3) refuging without foraging at night and foraging in the freestream during daytime (hereafter defined as a “combined” strategy). Based on Charnov’s diet model (1976), we assume that fish preferentially consume larger prey whenever available. Note that our approach does not take into account behaviours such as group hierarchy or predator avoidance and territoriality.

Finally, to translate model results of energetic gain to a value with biological meaning, we calculated a relative maximum benefit as

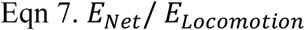

In this case, 100% relative maximum benefit indicates that a particular foraging strategy used for one day can result in one extra day of surplus energy before more food must be acquired, thereby designating energy available for other critical activities (e.g. migrating, growth, reproduction). Consequently, relative benefit provides an indication of the long-term benefit of adopting a particular feeding strategy in a way that reporting oxygen consumption units does not.

## Results

### Cost of swimming

Oxygen consumption increases with swimming speed for both refuging and freestream swimming fish. However, refuging fish did not increase their oxygen consumption until they swam faster than 68 cm s^-1^ (y = 66.6 + 0.30 * 1.07^x^, where y = oxygen consumption and x = swimming speed, repeated measures ANOVA, F_1,126_ = 72.9, p < 0.01). By comparison, fish in the freestream showed significantly greater oxygen consumption at each incremental swimming speed (y = 47.30 + 17.79 * 1.03^x^, Fig. 1).

**Fig. 1.**
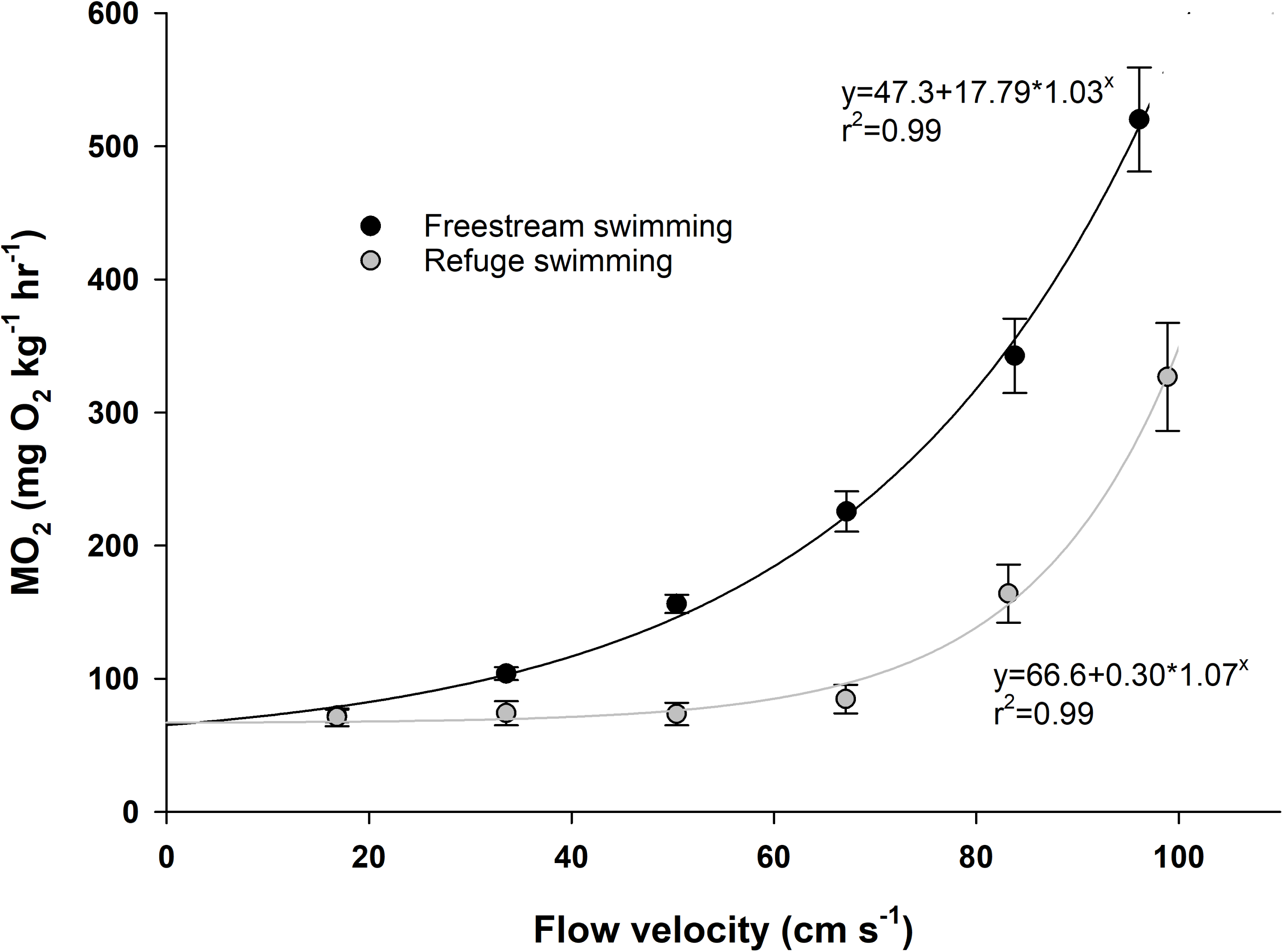
The energetic costs of swimming and refuging. Oxygen consumption of rainbow trout swimming in freestream flow (black line) compared to refuging behind a 5 cm diameter D cylinder (gray line). As flow velocity increases, the cost of swimming in the freestream increases more quickly compared to refuging behind a cylinder. The cost of refuging is similar across flow velocities until it exceeds 68 cm s^-1^. Error bars represent the s.e.m.

### Cost of prey attack

Oxygen consumption during prey attacks increased more quickly for refuging than for freestream swimming individuals (regression p ≤ 0.0001), indicating that the cost of each attack was greater for refuging trout (1.28 ± 0.18 mg O_2_ kg^-1^) than for trout in the freestream flow (0.78 ± 0.20 mg O_2_ kg^-1^, mean ± s.e.m., Fig. 2). For example, at a routine swimming speed of 68 cm s^-1^, foraging in the freestream was 40% less costly than compared to foraging while refuging (ANCOVA, F_1,31_ = 126.6, p < 0.01). This was speed dependent, for at a lower swimming speed of 10 cm s^-1^, this difference reduces to 6%.

**Fig. 2.**
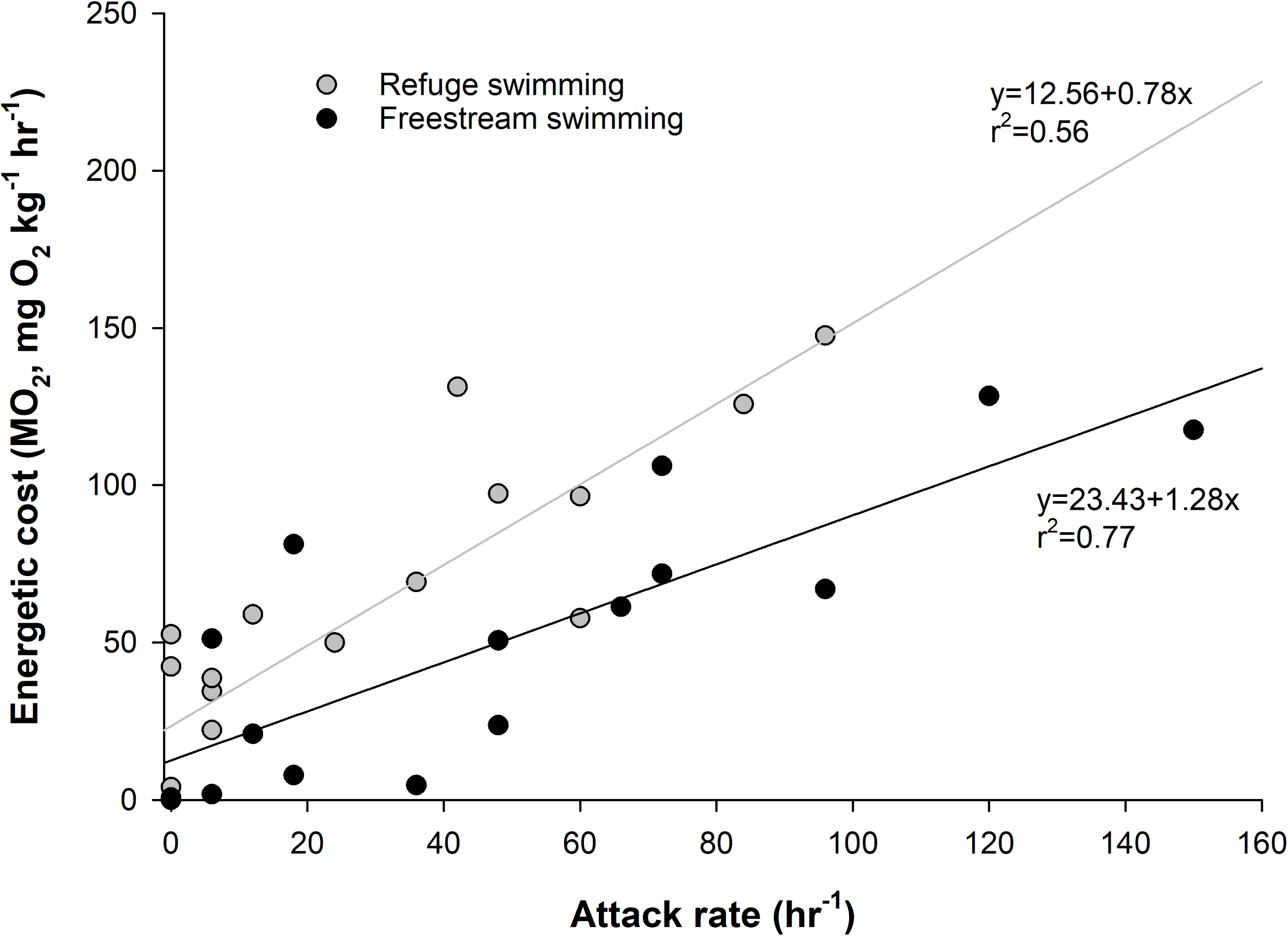
The energetic costs of attacking prey in flow. At a chosen flow velocity of 68 cm s^-1^, oxygen consumption of trout feeding on drifting prey increases linearly with attack rate during both freestream swimming and refuging (black and gray lines respectively). The slopes of the regression lines depict the energetic cost associated with each individual prey attack. The cost of attack is 64% greater during refuging compared to freestream swimming, reflecting the higher cost required to transition across a velocity gradient from refuging in a vortex street to intercepting prey in the freestream flow.

### Prey detection

Drift-feeding fishes must quickly detect and approach prey before they can capture it (Jenkins & Keeley 2010, Watz & Piccolo 2011). Both refuging and freestream swimming trout made the same number of attacks at prey (two-way ANOVA, F_1,58_ = 0.72, p = 0.58; F_4,58_ = 1.03, p_refuge_ = 0.32), indicating that the cylinder did not obscure prey detection for refuging trout. The proportion of prey attacked decreased as flow velocity increased for both behaviors (F_4,58_ = 21.05, p < 0.01; Fig. 3A).

**Fig. 3.**
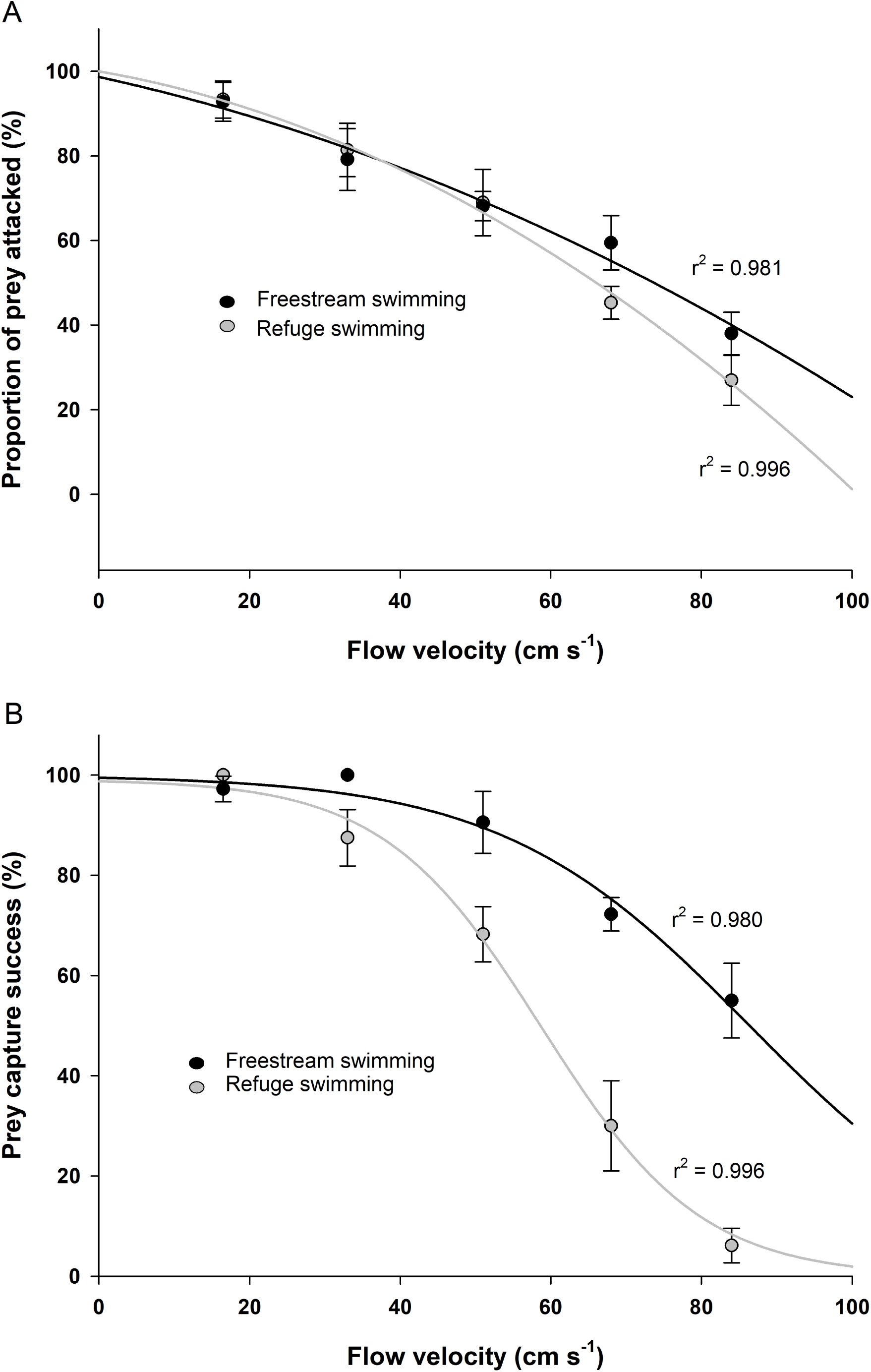
Attack rate and capture success for drift-feeding trout across flow velocities. (A) The proportion of prey attacked by trout swimming in the freestream versus refuging is not significantly different (black and gray lines respectively). In low flows (< 68 cm s^-1^), trout attack prey the majority of the time regardless of refuging behavior, while at higher flows (≥ 68 cm s^-1^) attack rate is comparatively lower in refuging trout. (B) As flow velocity increases, prey capture success decreases for both behaviors, but more substantially for refuging trout. Error bars represent the s.e.m.

### Prey capture success

Prey capture success was lower for refuging individuals compared to swimming in the freestream (two-way ANOVA, F_1,63_ = 28.2, p < 0.01). This was most evident at the highest flow velocities ≥ 51 cm s^-1^. The success of both feeding strategies decreased with increased flow velocity (F_4,63_ = 45.8, p < 0.01). For a given success percentage, individuals in the freestream can inhabit faster flow environments than refuging individuals (Fig. 3B, Table S1).

### Data-driven Cost-Benefit Model outcome

Figure 4A shows the net energetic benefit for trout feeding continuously either while refuging (gray line) or in freestream flow (black line). At flows < 25 cm s^-1^, both foraging strategies are energetically identical. When flow velocities exceed 25 cm s^-1^ it is more energetically favorable for trout to feed in freestream flow. The model predicted best 24 hr strategy is to combine freestream feeding and refuge swimming, where fish feed in freestream flow for 12 hrs and then refuge (without feeding) for 12 hrs (i.e. “combined” strategy, dashed line, Fig. 4A).

**Fig. 4.**
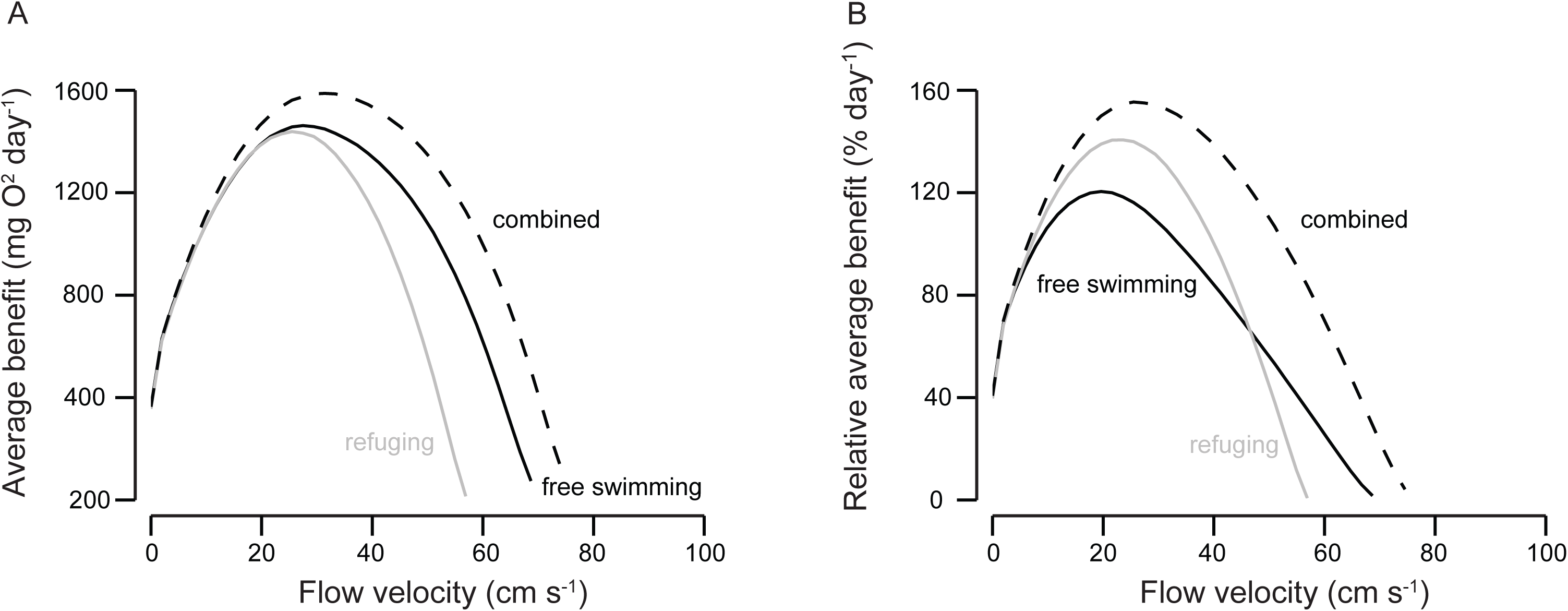
A model of the average absolute (A) and relative (B) energetic benefit for different feeding strategies in flow. (A) For flow velocities of < 25 cm s^-1^, the absolute benefit (defined as the energetic gain from food acquisition minus the cost of acquisition) of foraging exclusively in either freestream flow or behind a refuge is similar (black and gray line respectively). At flow velocities 25 - 70 cm s^-1^, it is energetically more beneficial to forage in the freestream flow as compared to refuging. (B) By dividing the absolute benefit with the cost of locomotion the relative benefit of each foraging strategy can be assessed (a value of 100% is defined as a full day surplus energy). At flows 10 – 50 cm s^-1^, foraging while refuging provides the greatest relative benefit due to the low difference in swimming cost between the refuge and the freestream. At swimming speeds greater than 50 cm s^-1^, however, foraging in the freestream is the energetically most advantageous strategy because the lower cost of foraging in the freestream outweighs the added cost of attack from a refuge. The highest energetic benefit over a 24hr period is realized by refuging without feeding for 12 hrs and foraging in the freestream for 12 hrs at all flow velocities > 20 cm s^-1^ (stippled line). This combined strategy also enables an energetic benefit to be gained from a greater range of flow velocities than possible for either behavior alone (up to 75 cm s^-1^).

Under average conditions, refuging trout swimming at 25 cm s^-1^ can acquire enough energy in a single day to last an additional ∼ 1.4 days (140%). This value is ∼ 1.2 days for fish foraging in the freestream and ∼ 1.5 days for trout that adopt the combined strategy defined above (dashed line, Fig. 4B).

We found that optimal feeding behavior depends on the magnitude of the flow velocity. For flows less than 50 cm s^-1^, refuging provides a greater relative energetic benefit than freestream swimming. For flows greater than 50 cm s^-1^, the benefit of refuging disappears and it is better to forage continuously in the freestream. However, the best overall strategy is to forage half the time in freestream flow and to refuge without feeding for the remaining time (dashed line). This strategy greatly expands the range of flow velocities where fish can maintain an energetic surplus, up to 75 cm s^-1^ in our model (Fig. 4B).

### Prey size and density requirement

To compensate for the increasing energetic costs of foraging, trout must feed on increasingly larger and more abundant prey as flow velocity increases (Fig. 5, S1). Prey size and density requirements increase faster with flow velocity for refuge-feeding individuals compared to freestream-feeding individuals. As a result, refuging individuals would in theory require 8 mm long prey when swimming at 70 cm s^-1^ whereas freestream individuals only need these prey sizes at 90 cm s^-1^ (Fig. 5A). Similarly, refuging individuals require densities of at least 15 prey per m^3^ of water volume in a flow of 60 cm s^-1^, whereas freestream individuals only need a density of 3 prey per m^3^ of water volume at the same flow velocity (Fig. S1). When fish combine both refuging and freestream swimming behaviors, their locomotion costs are minimal, which allows them to eat smaller and fewer prey than if they were exclusively refuging or swimming in freestream flow (Fig. 5B) and occupy flows with less prey (Fig. S1). Regardless of foraging strategy our model predicts that trout need to feed on prey larger than 2 mm, at density patches of at least 2 prey per m^3^, and conduct a minimum of 20 attacks per hour in order to obtain enough energy to sustain foraging costs (Fig. 5).

**Fig. 5.**
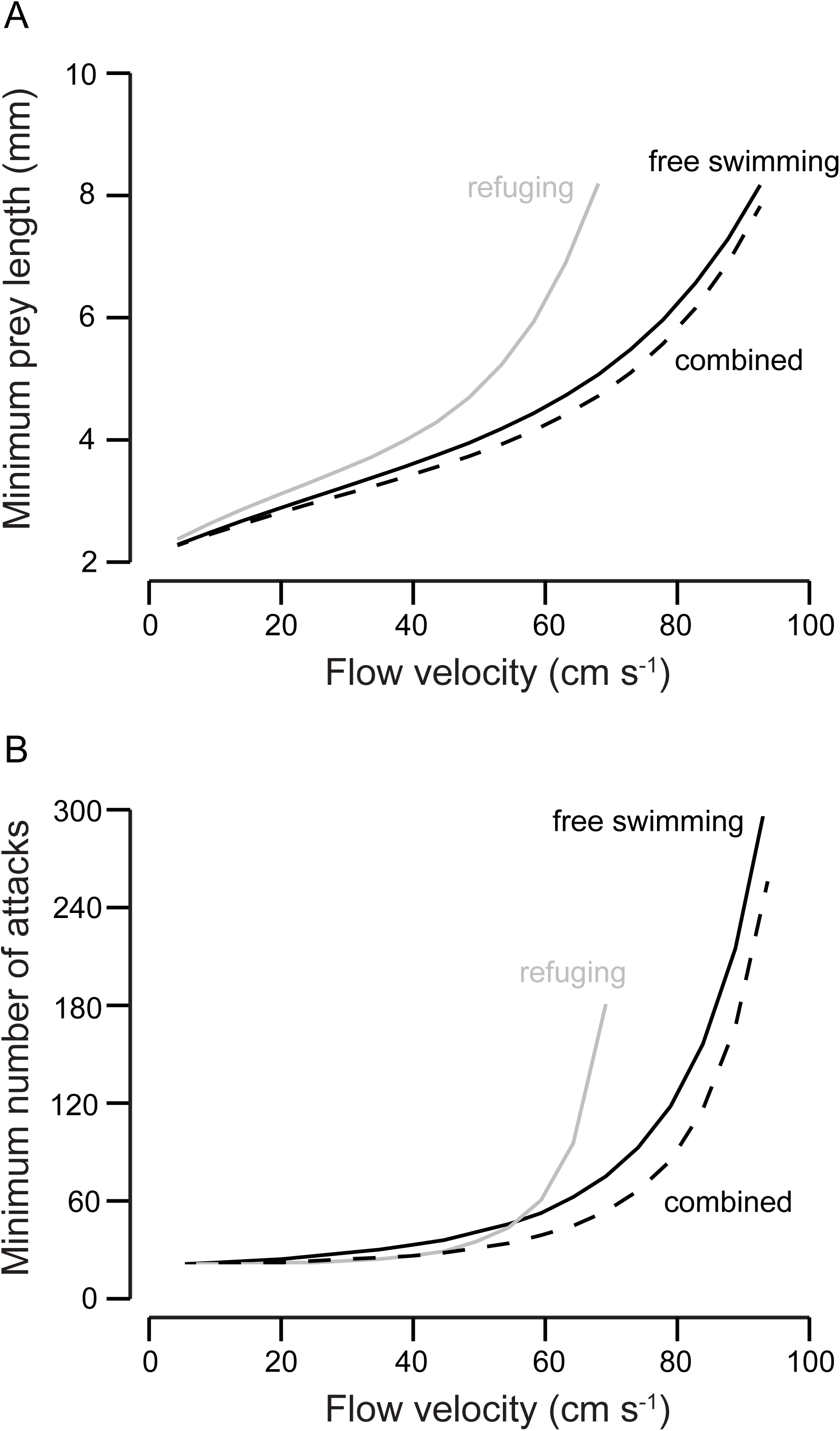
A model of the preferred prey size for drift-feeding trout at a given flow velocity and the minimum number of attacks required to provide an energetic surplus. (A) Taking into account the caloric content of the prey, the cost of locomotion and attack while refuging behind a cylinder or swimming in the freestream, and the likelihood of prey capture, trout should not target prey less than 2 mm in length. Relative to free swimming (black line) at a given flow velocity, trout that are solely refuging behind a cylinder (gray line) need to capture larger, higher calorie content prey due to the added attack costs originating from a refuge and reduced likelihood of capture. (B) Minimum number of attacks on prey required for a locomotion strategy to be energetically favorable. Minimum attack numbers at each flow velocity are defined as the lowest value of the following parameters: maximal successful prey captures per hour, number of prey available per hour, and maximum food intake per hour based on prey size availability. A foraging strategy consisting of feeding in the freestream and refuging while not feeding (stippled line) allows trout to minimize prey attack numbers across all flow velocities above 50 cm s^-1^.

## Discussion

Understanding what drives species distributions in nature is one of the fundamental goals of ecology. The energetics of feeding and locomotion are widely acknowledged to play a major role in influencing the distribution, abundance and behavior of organisms, but direct measurements of their costs are often lacking (Hughes & Kelly 1996, Grossman 2014, Piccolo et al 2014). In many aquatic ecosystems, fishes preferentially occupy high flow habitats (Guensch et al 2001, Fulton & Bellwood 2005, Grossman 2014, Johansen 2014, Piccolo et al 2014), despite the high associated costs of locomotion (Fulton et al 2013). It has long been assumed that fishes in high flows should seek to exploit refuges to reduce foraging costs (Liao et al 2003a, Johansen et al 2007, Johansen et al 2008, Taguchi & Liao 2011, Piccolo et al 2014, Rosenfeld et al 2014). However, our empirical cost-benefit model challenges this assumption by showing the opposite: in high flow, it is most energetically favorable for fish to either 1) refuge without foraging or 2) forage in the freestream without refuging. These results are primarily due to the cost of each attack relative to attack success rate, and may help to explain why trout often forage heavily in relatively open, freestream sections of rivers during an insect hatch (Grossman & Boule 1991). At the highest flow velocity tested (100 cm s^-1^), we found that all foraging strategies fail, even though trout both in our experiments (Fig. 1) and in nature (Cocherell et al 2011) are physically capable of swimming at these speeds and beyond. This is because once foraging costs are considered, swimming at these speeds is not energetically favorable, which may explain why fish are not typically found in such high flows in nature, and will vacate habitats once flow velocities exceed certain levels (Gido et al 2000, Cocherell et al 2011).

The capacity to exploit refuges to save locomotor energy has frequently been used to explain fish distributions in high flow habitats such as rivers and exposed coral reefs, where access and monitoring may be difficult (Hughes & Dill 1990, Liao et al 2003a, Johansen et al 2007, 2008, Urabe et al 2010, Taguchi & Liao 2011). This study confirmed this capacity in trout, revealing an almost 50% reduction in swimming costs at high speeds (> 65 cm s^-1^). However, we also observed a large (∼ 65%) and unexpected increase in the cost of attacking prey in the freestream while refuging, presumably due to the high cost of traversing a strong velocity gradient when leaving a vortex street refuge (Liao et al 2003b). Such a significant increase in energy usage for feeding has only previously been theorized (Puchett & Dill 1984, Boisclair & Tang 1993, Hughes & Kelly 1996, Guensch et al 2001, Rosenfeld & Boss 2001, Jenkins & Keeley 2010). The energetics of foraging have been identified as one of the important drivers of animal movement (Pyke 1984). Here, we directly reveal the mechanism underlying the substantial increase in energy use during foraging for the first time in drift-feeding fishes.

Fishes should seek to minimize the combined cost of metabolic maintenance, swimming and prey attack relative to the energetic gains from each attack. The optimal strategy likely depends on flow velocity and available prey size. By including our data on the energetic cost of attack into the total daily energy budget, we can derive the specific range of conditions under which it may be more energetically beneficial to exploit or avoid a flow refuge. Previous studies indicate that fishes interact with flow refuges using at least three different strategies: 1) reside in a refuge and only enter the freestream during an attack 2) reside and forage in the freestream, or 3) forage in the freestream and seek refuge when not foraging (Hill 1989, Grossman & Boule 1991, Guensch et al 2001, Piccolo et al 2014). When we use our data-driven model to explore these three strategies we find that there is an optimum range of flow velocities (within approximately 10 - 50 cm s^-1^, Fig. 4) that provides the greatest energetic benefit to trout. It is perhaps not surprising that this velocity range is equivalent to the natural conditions that are typically observed for rivers that contain the greatest abundance of trout (Hill & Grossman 1993, Guensch et al 2001, Urabe et al 2010).

Why, in low flow environments, is foraging while refuging better than foraging in the freestream flow? We show that when trout forage in currents less than 50 cm s^-1^, they can on average acquire 1.4 times their daily energy requirements (and even higher under optimal conditions). This energy surplus is possible due to the low cost of locomotion behind the refuge. In addition, slower currents lead to a reduced velocity gradient between the refuge and the freestream flow, which minimizes the cost of exiting the vortex street to attack prey. The lower cost of attack and higher attack success rate in slower currents allows trout to acquire surplus energy, which can then be used for growth, maintenance and reproduction.

In high flow habitats (> 50 cm s^-1^), however, the benefit of refuging disappears. We found that refuging trout suffered a net energetic loss due to a 65% increase in the cost of attacking prey and a 40% reduction in attack success rate, demonstrating that it is costlier to forage from a refuge than from freestream flow. Specifically, the energetic benefit of refuging while foraging is inversely related to flow velocity. We suggest that this is because the energy needed to accelerate from a place of refuge into faster freestream flow necessitates traversing across a steep velocity gradient. Compared to foraging in freestream flows, refuging fish that dart out to capture food are expected to accelerate more quickly in order to overcome these velocity gradients. The large amplitude kinematics of fast accelerations is significantly different than those of steady swimming (Akanyeti et al 2017), and is likely the main contributor to the high energetic requirements of this behavior. This leads to an inability of feeding trout to maintain an energy surplus while refuging because the cost of each attack is greater than energy content of the prey. Contrary to prevailing theory, our model predicts that individuals living in flows > 50 cm s^-1^ should completely avoid refuges while foraging. This counterintuitive insight is only possible because we employ the first direct measurements of feeding costs in controlled conditions. Our results explain field studies in which trout have been reported to forage predominantly in the freestream sections of rivers (Grossman & Boule 1991). At the highest flow velocities in our study (> 81 cm s^-1^), trout can only ensure adequate energy intake by foraging in the freestream when prey densities are greater than 15 individuals per m^3^ (Fig. S1.). This suggests that trout in rivers will not occupy regions of freestream flow during periods of low prey density.

Our results highlight the importance of considering the relatively high cost of foraging when refuging, which has been absent in previous energetic models (see Piccolo et al 2014 for review). Perhaps most importantly for fisheries conservation managers, we argue that a classification of optimum habitat conditions cannot be determined solely on the basis of swimming energetics. Our data suggests that preferred flow velocities in the field are limited not only by swimming capacity, but also by the energetic demands of foraging. This observation may also help explain potential discrepancies between predicted and realized habitat and movement patterns in the wild.

Our results lead us to suggest that the main benefit of feeding from a refuge may not be to save in locomotion costs (which occurs only at lower flow velocities), but as a strategy for resilience to unpredictable food conditions. Temporal fluctuations in food availability are common in nature (Nakano et al 1999, Hamner et al 2007, Jenkins & Keeley 2010, Armstrong & Schindler 2011). In unpredictable habitats, individuals that occupy flow refuges will require less energy over time and will be more resilient to prolonged periods of low food availability. Such refuging patterns have been observed in rivers that experience frequent periods of low prey abundance (Hughes & Dill 1990) and pulses of high current (Cocherell et al 2011). Within an optimal range of flow velocities (here 10 – 50 cm s^-1^), refuging individuals also require less calorie rich prey to sustain the same energetic benefits as individuals swimming in freestream flow. This allows refuging individuals to forage on a wider variety of prey that may be smaller and have less caloric content.

As flow velocity increases, the cost of swimming and foraging also increases while the ability to detect and capture prey decreases (Hill & Grossman 1993, Braaten et al 1997). Trout may deal with this problem by only targeting larger, calorie-rich prey in order to compensate for the diminished opportunities to capture prey. In nature, invertebrate prey of 8mm or greater make up less than 1% of all drifting food (Table 1), yet trout preferentially select these larger prey (Nakano et al 1999). It must be noted, however, that faster flows generally delivery more and larger prey over a unit of time (Hill & Grossman 1993, Hayes et al 2007, Jenkins & Keeley 2010). Based on prey size distribution and caloric prey content in natural rivers, our energetic model predicts the preferred prey size of foraging trout under different feeding strategies and flow velocities. We predict that refuging trout occupying habitats with flows greater than 65 cm s^-1^ should solely select prey sizes of 8 mm or greater in order to account for the costs of foraging. The need to target large prey based on energetics may explain why trout have been observed to leave sections of rivers that appear to contain abundant, but small prey (Hughes & Dill 1990, Gido et al 2000). Likewise, drift-feeding trout in fast flows appear to avoid small prey even when their abundance is greater by several orders of magnitude than larger prey (Hill & Grossman 1993, Nakano et al 1999).

Our work suggests that, regardless of flow velocity, the minimum prey size that trout should eat must be 2 mm long in order to provide enough energy (equivalent to ∼2.1 joule) to cover the energetic cost of a successful capture (Fig. 5). The majority of invertebrate prey found in rivers ranges from 0.1 - 10.0 mm in length, with more than 50% of all prey 2.0 mm or less (Table 1). Therefore, it may not be energetically favorable for trout to forage on the majority of available prey. Indeed, examinations of stomach contents have revealed that trout preferentially forage on prey greater than 2.0 mm in length (Hill & Grossman 1993, Braaten et al 1997, Nakano et al 1999, Guensch et al 2001, Hughes et al 2003). This discrepancy between prey size availability and stomach content analyses has previously been attributed to large gill raker spacing, which may not be effective for filtering and retaining smaller prey (Bisson 1978, Hughes et al 2003). We do not believe this to be the case, given that trout are not filter feeders but target and capture individual prey. In addition, true filter feeding fishes can capture prey much smaller than their gill raker spacing due to a unique vortex filtration mechanism (Motta et al. 2010, Paig-Tran et al. 2013). It is not known whether trout possess such a filtration mechanism. Here, we provide an alternative explanation based on energetics. Fishes foraging on individual prey in flow such as trout and certain coral reef fishes should consider the energy content of each prey, which over time must be greater than the cost of its acquisition. To our knowledge, this is the first direct experimental demonstration that supports an energetics argument to explain such selective foraging patterns in fishes.

Values of foraging costs in flow should be considered for cost-benefit models, along with consideration for prey detection and capture abilities, in order to improve their accuracy from existing models in which these parameters are lacking (Hughes and Dill 1990, Hughes and Kelly 1996, Guensch et al 2001, Hughes et al 2003, Urabe et al 2010, Piccolo et al 2014). We account for these parameters here, leading us to believe that our general approach can be broadly applied to both freshwater and marine species, though the details of prey size and energetic costs may differ. The utility of our approach is that it predicts energetically favourable flow velocities for fishes (herein described as optimum flow range). As such, it provides a mechanistic basis for understanding why individuals in nature are typically found associating with refuges only within a specific range of flow velocities. This approach, where links between energetic demands and habitat usage are demonstrated, is starting to illuminate our understanding of distribution patterns in a variety of species (Urabe et al 2010, Rosenfeld et al 2014). The ability to determine *a-priori* the optimum flow requirements will help better predict movement and habitat usage patterns not only for salmonids in rivers, but of other ecologically and commercially important species (e.g. Kiflawi & Genin 1997, Zeller 2002, Johansen et al 2014, 2015). As such, we believe that our model has strong conservation implications that can be applied broadly.

By using respirometry to directly measure foraging costs in the lab, our empirical approach provides critical new insight that challenges established assumptions of ecology and behavior in current swept ecosystems. We demonstrate that in high flow habitats, hydrodynamic refuges are not energetically favorable locations, and that the best feeding strategy across flow velocities is adaptive; refuging in slower flows when not foraging and only foraging in faster freestream flows. Our experimental results provide a framework to understand the mechanisms underlying habitat preferences and movement patterns in current-swept environments, and generates hypotheses that can be tested to see how well these strategies are employed by fishes in nature;

Conceptualization: J.L.J., J.C.L; Methodology: J.L.J., J.C.L; Software: O.A., J.C.L; Validation: O.A., J.L.J.; Formal analysis: O.A.,J.L.J.; Investigation: J.L.J., J.C.L; Resources: J.C.L; Data curation: J.L.J., J.C.L; Writing - original draft: J.L.J.; Writing - review & editing: O.A.,J.L.J., J.C.L; Supervision: J.C.L.; Project administration: J.C.L; Funding acquisition: J.C.L.

## Acknowledgements

We would like to thank Ashley Peterson for fish care and Masashige Taguchi for help in collecting data and preliminary analyses. All protocols were approved by the University of Florida Institutional Animal Care and Use Committee. This research was supported by the National Institute on Deafness and Other Communication Disorders Grant RO1-DC-010809 and National Science Foundation Grants IOS 1257150 and IOS 1856237 to J.C.L. The authors declare no competing interests.

## Supplementary data

**Table S1.** Post Hoc Planned comparison of prey capture success against control (16 cm s^-1^ flow velocity). Significance accepted at P_cutoff_ < 0.01.

|  | Flow velocity (cm s<sup>-1</sup>) | t | p |
| --- | --- | --- | --- |
| Free stream | 33 | 0.644 | 0.52 |
|  | 51 | 1.019 | 0.31 |
|  | 68 | 4.350 | <0.01 |
|  | 84 | 5.776 | <0.01 |
| Refuging | 33 | 2.001 | 0.05 |
|  | 51 | 4.308 | <0.01 |
|  | 68 | 6.938 | <0.01 |
|  | 84 | 8.485 | <0.01 |

**Fig. S1**. Minimum prey density required to gain an energetic surplus across flow velocity. At flows < 10 cm s^-1^, a high prey density is required due to the low delivery rate of prey. Within flow velocities of 10 – 50 cm s^-1^, required prey density falls below 2 prey m^-3^ due to the lower cost of attack and increased prey capture success. At higher flow velocities, minimal prey density increases rapidly due to lower capture success rate and increased cost of attack. Note that refuging individuals (gray line) require the highest prey density when flow velocities exceed 50 cm s^-1^ due to the greater cost of attack.

